# Catching a liar through facial expression of fear

**DOI:** 10.1101/2021.02.24.432646

**Authors:** Xunbing Shen, Gaojie Fan, Caoyuan Niu, Zhencai Chen

**Affiliations:** Jiangxi University of Traditional Chinese Medicine, Nanchang, 330004; Louisiana State University, Baton Rouge, 70803

**Keywords:** Deception detection, leakage theory, fear, machine learning, asymmetry

## Abstract

The leakage theory in the field of deception detection predicted that liars could not repress the leaked felt emotions (e.g., the fear or delight); and people who were lying would feel fear (to be discovered), especially under the high-stake situations. Therefore, we assumed that the aim of revealing deceits could be reached via analyzing the facial expression of fear. Detecting and analyzing the subtle leaked fear facial expressions is a challenging task for laypeople. It is, however, a relatively easy job for computer vision and machine learning. To test the hypothesis, we analyzed video clips from a game show “The moment of truth” by using OpenFace (for outputting the Action Units of fear and face landmarks) and WEKA (for classifying the video clips in which the players was lying or telling the truth). The results showed that some algorithms could achieve an accuracy of greater than 80% merely using AUs of fear. Besides, the total durations of AU 20 of fear were found to be shorter under the lying condition than under the truth-telling condition. Further analysis found the cause why durations of fear were shorter was that the duration from peak to offset of AU20 under the lying condition was less than that under the truth-telling condition. The results also showed that the facial movements around the eyes were more asymmetrical while people telling lies. All the results suggested that there do exist facial clues to deception, and fear could be a cue for distinguishing liars from truth-tellers.

## 1. Introduction

Are there any observable behaviors or cues which can differentiate lying from being honest? For this question, almost all researchers in the field of deception detection think there is no “Pinocchio’s nose”(DePaulo et al., 2003). Nevertheless, Many researchers try hard to find the cues to deception (Denault et al., 2020; Levine, 2018). Specifically, from the perspective of leakage theory (Ekman, 2003; Ekman & Friesen, 1969; Matsumoto & Hwang, 2020; Porter et al., 2011; Porter et al., 2012; Su & Levine, 2016), observable emotional facial expressions (microexpressions and macroexpressions) can, to some degree, determine who is lying and who is telling the truth (It’s a probability problem, see Levine, 2018, 2019).

The “leakage theory” asserts that high-stake lies (the rewards come with serious consequences or there can be severe punishments) can result in ‘leakage’ of the deception into physiological changes or behaviors (especially microexpressions). In turn, the presence of leakage suggests the high probability of existence of deception(Ten Brinke, MacDonald, et al., 2012; ten Brinke & Porter, 2012; Ten Brinke, Porter, et al., 2012). However, there is debate about whether or not the emotional facial expressions can differentiate lying from truth-telling. Some researchers (Matsumoto & Hwang, 2018; ten Brinke & Porter, 2012; Ten Brinke, Porter, et al., 2012) thought the emotional facial microexpression could be a cue to lies and found some evidence supporting the claim. Nevertheless, Burgoon (2018) regarded that the microexpressions were not the best way to catch a liar. Furthermore, Vrij et al. (2019) even categorized microexpression into pseudoscience.

There indeed are some behavioral cues that can, to some degree, differentiate lying from truth-telling(Vrij et al., 2006; Vrij et al., 2000). Especially, pupil dilation and pitch are closely related to lying (Levine, 2018, 2019). Emotional facial expressions can also be behavioral cues of this kind. Most of the deception researchers agree that lying does involve processes or factors such as arousal and felt emotion (Zuckerman et al., 1981). Meanwhile, there are involuntary aspects of emotional expression. As noted by Darwin, some actions of facial muscles are the most difficult to control voluntarily and are the hardest to be inhibited (the so-called Inhibition Hypothesis, see also (Ekman, 2003). When a strong felt genuine emotion presents, the actions of the expressions of the felt emotion cannot be suppressed (Baker et al., 2016). Hurley and Frank (2011) provided evidence for Darwin’s hypothesis and found that deceivers could not control some elements of their facial expression, such as eyebrow movements. The liar would feel fear, duping delight, disgust, or appear tense while lying, and would attempt to suppress these emotions by neutralizing, masking, or simulating (Porter & Ten Brinke, 2008). However, the liars cannot inhibit them completely and the felt emotion will be “leaked” out in the form of microexpressions, especially under high-stake situations (Ekman & Friesen, 1969).

Some recent research substantiated the claim of emotional leakage (Porter et al., 2011; Porter et al., 2012). When liars camouflage with an unfelt emotional facial expression or neutralize the felt emotion, at least one inconsistent expression would leak and present transiently (Porter and Ten Brinke (2008). ten Brinke and Porter (2012) showed that liars would present unsuccessful emotional masking and certain leaked facial expressions (e.g., “the presence of a smirk”). In addition, they found that false remorse was associated with (involuntary and inconsistent) facial expressions of happiness and disgust (Ten Brinke, MacDonald, et al., 2012).

There is some evidence that supports the claim that leaked emotions can differentiate telling lies from telling the truth. Wright Whelan et al. (2014) used cues that included emotional ones to identify high-stake deception and got an accuracy of 78%. Meanwhile, Wright Whelan et al. (2015) found non-police and police observers could reach an accuracy of 68% and 72%, respectively. Using methods of machine learning, Su and Levine (2016) found that emotional facial expressions (including microexpressions) could be effective cues while the participants judging high-stake lies, in which the accuracy was much higher than those reported in previous studies (e.g., Bond Jr & DePaulo, 2006). They found Action Units (the contraction or relaxation of one or more muscles, see Ekman & Friesen, 1976) of AU1, AU2, AU4, AU12, AU15, and AU45 (blink) could be potentially effective indicators for distinguishing liars from truth-tellers in high-stakes situations. Bartlett et al. (2014) showed that machine vision could differentiate deceptive pain facial signals from genuine pain facial signals (at 85% accuracy). Matsumoto and Hwang (2018) found that facial expressions of negative emotions that occurred for less than 0.40 and 0.50 seconds could differentiate truth-tellers and liars.

The leakage theory of deception predicted that liars should fear of being discovered, and that the fear emotions resulted from deception (especially high-stake one) might leak the deception (Levine, 2019). Meanwhile, it is presumed that if the fear associated with deception is leaked, then the duration of the leaked fear will be shorter due to the nature of leaking (which will be showed as fleeting fear micro-expressions) and repressing. Someone may argue that the fear emotion may also appear while telling the truth. It can be true. Nevertheless, for a truth-teller, the fear of being wrongly treated as a liar would be less leaking, since a truth-teller doesn’t need to try hard to repress the fear as liars do (the degree of repressing will be different between liars and truth-tellers). On average, the duration of fear (or AUs of fear) in lying situations would be shorter than that in truth-telling situations due to the harder repressing. Meanwhile, researchers (Ekman et al., 1981; Frank et al., 1993) found that the genuine smile has different dynamic features, such as a smoother onset and more symmetry(Ekman et al., 1981), when compared with a deliberate smile. Accordingly, the leaked emotional facial expressions of fear while lying and the less leaked ones when telling a truth may have different dynamic qualities.

Stakes may play a vital role while using an emotional facial expression as a cue to deception. Participants experience fewer emotions or less cognitive load in laboratory research (Buckley, 2012). Almost all laboratory experiments are typical of low stakes and are not sufficiently motivating to trigger emotions giving rise to leakage (in the form of microexpressions). Consequently, liars in laboratory experiments are not as nervous as in real-life high-stake situations, with no or little emotion leakage. As noted by Vrij (2004), some laboratory-based studies in which the stakes were manipulated had found that high-stakes lies were easier to detect than low-stakes lies. Frank and Ekman (1997) stated that “*the presence of high stakes is central to liars feeling strong emotion when lying*”. Therefore, lying under the higher stakes condition would be more detectable while using cue of emotional facial expressions, and leaked emotional facial expressions may mostly occur in a high-stakes context.

Hartwig and colleagues (2014) claimed that the emotional leakage theory could not be supported and the context of the high stake would influence both liars and truth-tellers, as liars and truth-tellers might experience similar psychological processes. In other words, a truth-teller would also produce inconsistent emotional expressions like fear. To some degree, this is the case (ten Brinke & Porter, 2012). Even though the high-stake situations increase pressure on both liar and truth-tellers, it can be assumed that the degree of increment would be different; and the liars would feel much higher pressure. In addition, to fabricate a lie, in general, liars have to think more in their minds and would have higher emotional arousal than truth-tellers. Consequently, for liars, the frequency or probability of leaking an inconsistent emotional expression (say, fear) would be higher and there would be more emotional signs presented for liars. In theory, the higher the stakes are, the more likely cues associated with deception (e.g., fear) are leaked, and the easier the liars could be identified.

Based on the leakage theory and previous evidence, we hypothesize that 1) emotional facial expressions of fear (fear of being caught) can differentiate lying from truth-telling at high-stake situations; 2) The duration of AUs of fear in lying will be shorter than in truth-telling; 3) The symmetry of facial movements will be different, as facial movements in lying situations will be more asymmetrical (due to the nature of repressing and leaking).

To test these hypotheses, we used videos of high-stake lies as experimental material, and a software of computer vision to automatically analyze the signals of emotional facial expressions. Compared to the slightly-better-than-chance accuracy obtained by human observers, computer vision can reach a relatively high accuracy when distinguishing deception from truth-telling (Bartlett et al., 2014). Given that the subtle differences of emotional facial expressions may not be detected by naive human observers, the methods of computer vision may capture the different features between lying and truth-telling situations which cannot be perceived by a human lie detector.

## 2. Results

### 2.1 AUs of fear can differentiate liars from truth-tellers

#### 2.1.1 Machine learning classification results

The whole dataset was split into two subsets, i.e., data collected from 12 participants were used for training, and the data collected from remaining 4 participants were used for testing. Three classifiers were trained on dataset of 12 participants to discriminate liars from truth-tellers using feature vectors of AUs of fear (i.e., AU01, AU02, AU04, AU05, AU07, AU 20, and AU26, for details see https://imotions.com/blog/facial-action-coding-system/). All of the three classifiers, Random Forest, K-nearest neighbours (LBK), and Bagging, were trained in WEKA via a 10-fold cross-validation procedure. To highlight the relative importance of AUs of fear in classification accuracy, we eliminated all other indicators used by Beh and Goh (2019). Table 1 shows the performance of machine learning analysis which conducted on dataset of 12 participants and tested with the data of remaining 4 participants.

**Table 1.**
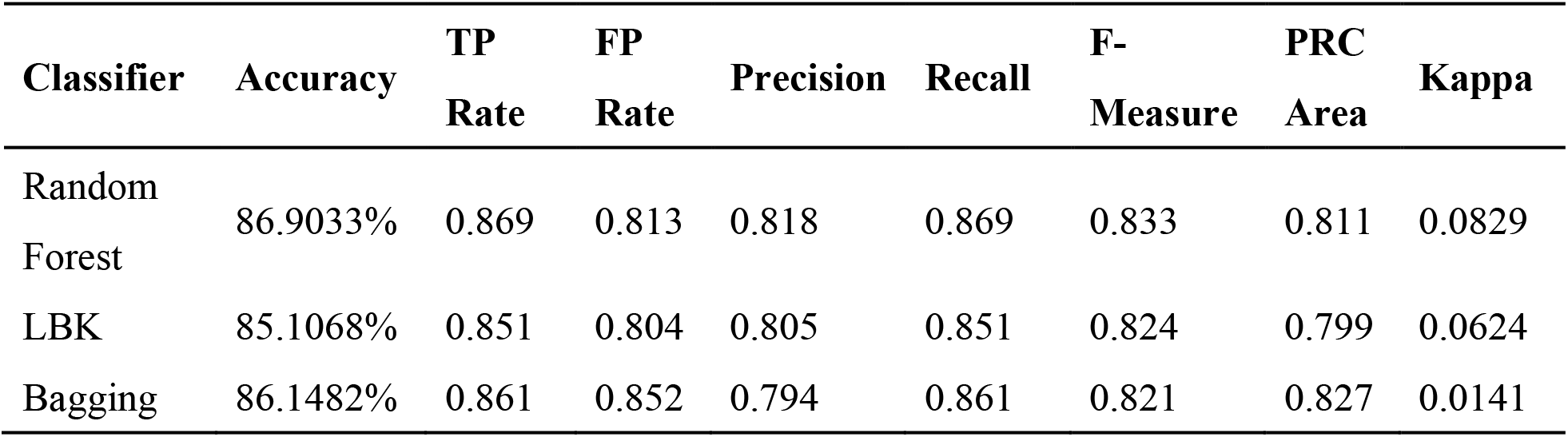
Machine learning performance of the Random Forest, LBK, and Bagging.

| Classifier | Accuracy | TP Rate | FP Rate | Precision | Recall | F-Measure | PRC Area | Kappa |
| --- | --- | --- | --- | --- | --- | --- | --- | --- |
| Random Forest | 86.9033% | 0.869 | 0.813 | 0.818 | 0.869 | 0.833 | 0.811 | 0.0829 |
| LBK | 85.1068% | 0.851 | 0.804 | 0.805 | 0.851 | 0.824 | 0.799 | 0.0624 |
| Bagging | 86.1482% | 0.861 | 0.852 | 0.794 | 0.861 | 0.821 | 0.827 | 0.0141 |

Table 1 reports the percentage of accuracy obtained on the testing data set. In addition to accuracies, the table reports the weighted average of True Positive Rate (TP Rate, instances correctly classified as a given class), False Positive Rate (FP Rate, instances falsely classified as a given class), Precision value (proportion of instances that are truly of a class divided by the total instances classified as that class), Recall value (proportion of instances classified as a given class divided by the actual total in that class), F-Measure (A combined measure for precision and recall), Precision-Recall Curve (PRC) Area value (A model performance metrics based on precision and recall) and Kappa (which measures the agreement between predicted and observed categorizations). The details of these statistics can be seen in Witten et al. (2016).

#### 2.1.2 the differences of AUs of fear between truth-telling and lying video clips

We took the averages of AUs related to fear for each individual to explore how they differ in lying versus truth-telling. The first analysis was carried out by examining the statistical differences of AUs of fear between truth-telling and lying video clips through paired *t*-test. To avoid the multiple-testing problem, we applied Bonferroni correction and set p-value to 0.007. We also calculated Cohen’s d to measure effect size. The results are presented in Table 2. When Bootstrapping was used, the p-value of comparing AU20 in the two groups was .006 (for AU05 the corresponding p-value is .008). This analysis revealed that liars and truth-teller have differences in the facial expressions of fear.

**Table 2.** the results of paired *t*-test for comparing the means of values of AUs of fear between truth-telling and lying video clips

| Feature | Deception<br>(Mean) | Truth<br>(Mean) | 95% CI of mean<br>difference |  | *t*-value | *p*-value | Effect<br>size* |
| --- | --- | --- | --- | --- | --- | --- | --- |
| AU01 | .2544 | .2735 | -.1562 | .1180 | -.297 | .771 | .074 |
| AU02 | .1308 | .1759 | -.1099 | .0196 | -1.487 | .158 | .371 |
| AU04 | .1686 | .1554 | -.0709 | .0972 | .333 | .743 | .084 |
| AU05 | .0341 | .0639 | -.0505 | -.0090 | -3.060 | .008 | .766 |
| AU07 | .7929 | .8517 | -.3581 | .2405 | -.419 | .681 | .105 |
| AU20 | .0838 | .1427 | -.0978 | -.0200 | -3.226 | **.006** | .807 |
| AU26 | .3969 | .4721 | -.1825 | .0321 | -1.493 | .156 | .374 |

\*Note: the effect sizes were calculated by using the calculator from the website: https://memory.psych.mun.ca/models/stats/effect_size.shtml.

| Feature | Deception (Mean) | Truth (Mean) | 95% CI of mean difference |  | *t*-value | *p*-value | Effect size* |
| --- | --- | --- | --- | --- | --- | --- | --- |
| AU01 | .2544 | .2735 | -.1562 | .1180 | -.297 | .771 | .074 |
| AU02 | .1308 | .1759 | -.1099 | .0196 | -1.487 | .158 | .371 |
| AU04 | .1686 | .1554 | -.0709 | .0972 | .333 | .743 | .084 |
| AU05 | .0341 | .0639 | -.0505 | -.0090 | -3.060 | .008 | .766 |
| AU07 | .7929 | .8517 | -.3581 | .2405 | -.419 | .681 | .105 |
| AU20 | .0838 | .1427 | -.0978 | -.0200 | -3.226 | **.006** | .807 |
| AU26 | .3969 | .4721 | -.1825 | .0321 | -1.493 | .156 | .374 |

\*Note: the effect sizes were calculated by using the calculator from the website: https://memory.psych.mun.ca/models/stats/effect_size.shtml.

### 2.2 There were more transient durations of AU of fear while lying

Ekman (2003) reported that many people could not inhibit the activity of the AU20 (Stretching the lips horizontally) while examining videotapes of people lying and telling the truth. Our results reported in section 2.1.2 also found significant differences between truth-telling and lying video clips in values of AU 20. Therefore, differences in the durations from onset to peak, from peak to offset, and total durations of AU 20 between truth-telling video clips (in which the quantity of AU20 is 675) and lying video clips (in which the quantity of AU20 is 47) were analyzed with independent samples *t*-test, using bootstrapping with 1000 iterations. The results showed that there were significant differences in the total duration and duration from peak to offset between truth-telling video clips and lying video clips (20.77 vs. 15.21 frames, *p* = .033, effect size = 0.276; 11.35 vs. 6.98 frames, *p* = .04, effect size =0.347). The durations of AU20 in lying video clips were nearly 4 frames (133 ms) shorter than those in truth-telling video clips on average, because the facial movements (herein the AU20) disappeared more quickly in the lying condition. Figure 1 shows the distribution of total frames, frames from onset to apex, and frames from apex to offset of AU20. The median is 12 in the truth-telling video clips and 8 in the lying video clips. For lying video clips, the 95% confidence interval is 10.32 to 20.11 frames for the mean of total duration, and 19.03 to 22.52 frames for truth-telling video clips. There were 16 (out of 47) AU20s which durations were less than or equal to 6 frames (200 ms) in the lying video clips, while there were 145 (out of 675) in the truth-telling video clips. There were 32 AU20s which durations were less than or equal to 15 frames (500 ms) in the lying video clips, and the corresponding number is 407 in the truth-telling video clips.

**Figure 1.**
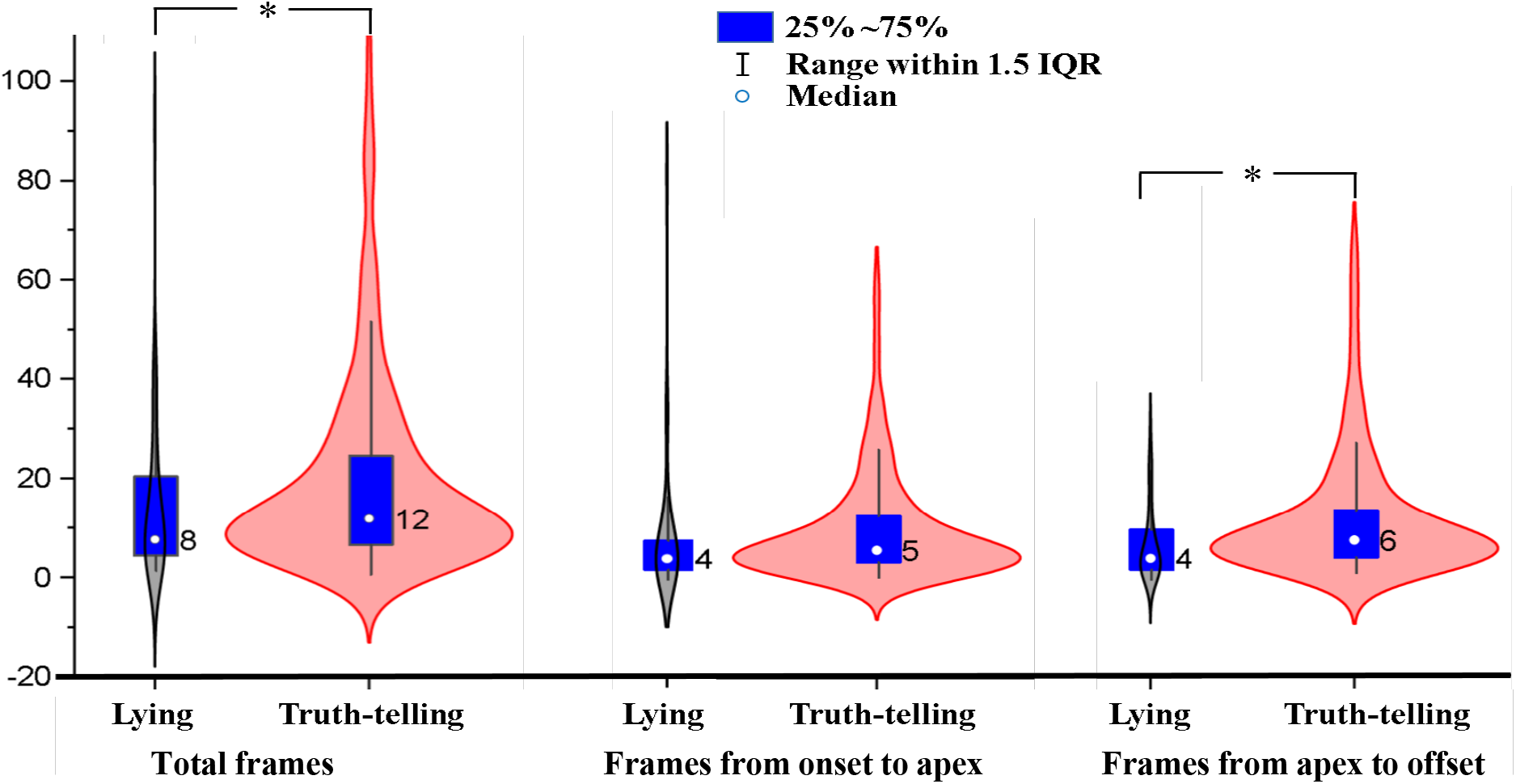
Violin plot for frames of AU20 in truth-telling and lying video clips. IQR = InterQuartile Range. *: statistically significant (p <.05) differences between lying and truth-telling.

### 2.3 Asymmetries of the facial movements were more salient in lying than truth-telling

Ekman et al. (1981) manually analyzed the facial asymmetry by using the Facial Action Coding System (FACS). This artificial approach is time-consuming, and subjective. In the current study, we proposed a method that used coherence (a measure of the correlation between two signals/variables) to measure the asymmetry. The more symmetrical the facial movements of the left and right face, the higher the coefficient of correlation between them. Consequently, the value of coherence (ranges from 0 to 1) can be a measurement of asymmetry or symmetry.

We calculated the distances of ld1 and rd1 (Beh & Goh, 2019) in each frame, which corresponded to movements of left and right eyebrows. Next, we used the MATLAB function of Wcohenrence (wavelet coherence) to measure the correlation between ld1 and rd1 in each video. If the movements were symmetrical, e.g., they have the exact same onset time, reach the apex on the same time, and disappear at the same time, the coherence between ld1 and rd1 should be 1, and any asynchrony would result in a value of coherence of less than 1, and the value of the coherence would be even smaller with the more asymmetry existed. Figure 2 shows the wavelet coherence in truth-telling and lying video clips.

**Figure 2.**
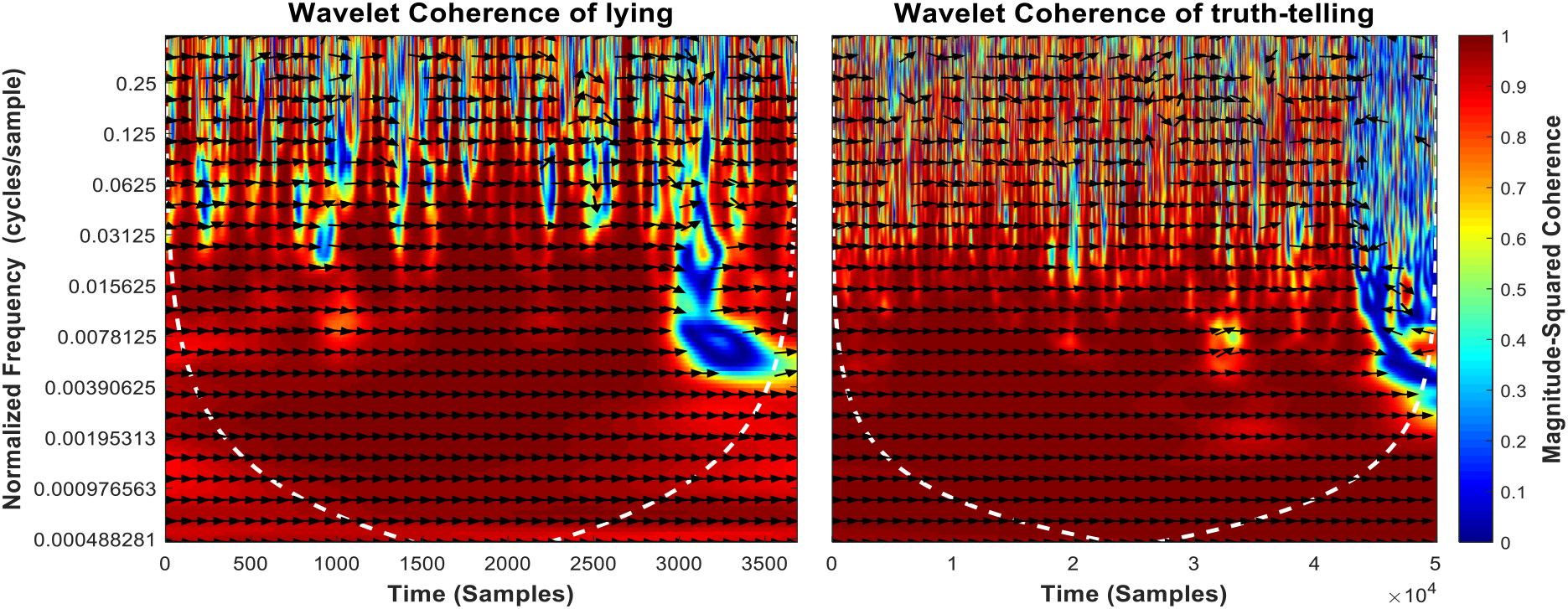
Squared wavelet coherence between the ld1 and rd1 in lying (left panel) and truth-telling (right panel) situations. The relative phase relationship is shown as arrows (A rightward arrow indicates 0 lag; a bottom-right arrow indicates a small lead of ld1; a leftward arrow indicates ld1 and ld2 is anti-correlated.).

The output values of the function of Wcohenrence for each player (i.e., the average of coherence between ld1 and rd1) were entered into the Permutation Test (see the following link for details: https://github.com/lrkrol/permutationTest) to compare the asymmetry differences between the lying and truth-telling situation. Permutation tests provide elegant ways to control for the overall Type I error and are distribution free. The results showed that there were significant differences between lying and truth-telling situations (the means of coherence are 0.7083 and 0.8096, *p* = .003, effect size = 1.3144).

## 3. Discussion

Is there any effective cue to deception? It is widely accepted that cues to deception, even exist, are weak. According to leakage theory, the leaked emotional facial expressions, especially the leaked fear, can differentiate lying from truth-telling. The current study confirmed the prediction of leakage theory.

The results of machine learning indicated that emotional facial expressions of fear can differentiate lying from truth-telling in the high-stake game show; the paired comparisons showed significant differences between lying and truth-telling in values of AU 20 of fear (AU5 is marginally significant). The results also substantiated the other two hypotheses. The duration of AUs of fear in lying was shorter than in truth-telling. The results showed that the total duration and the duration from peak to offset of AU 20 of fear were shorter while lying than while telling truth. The third hypothesis predicted that the symmetry of facial movements will be different, and the findings indicated that the facial movements were more asymmetrical in lying situations than in truth-telling situations.

In the current study, the method of machine learning can classify deception and honesty, which made up the shortcomings of human coding and were managed to find out the subtle differences between lying and truth-telling. Meanwhile, an objective measure of asymmetry was proposed. To our best knowledge, this is the first objective method to measure the asymmetry of facial movements in deception detection. By using these methods, we did find there were differences between lying and truth-telling, which is the prerequisite for looking for clues of deception (if there is no difference between lying and truth-telling, then there will be no cues to deception).

The leaked behaviors can be cues to deception, but they are not deception per se. They are, however, closely linked with deception. As shown in the results, truth-tellers also can experience fear. However, for honest people, the dynamics of experienced fear were very different when compared with liars. Thus, the fear emotion could be considered as a “hot spot” of deceit. Looking for the nonverbal “hot spots” of individuals is very suitable for the scenario in which rapid evaluation is required. Some other approaches of deception detection, for example, brain activities, cannot provide real-time results (Vrij & Fisher, 2020). The results suggested that the “hot spots” - emotional expressions of fear - could distinguish between truthful and deceptive messages with a reasonable level of accuracy. Using machine learning, we can get a relatively higher accuracy (above 80%) compared to the average accuracy achieved by people (54%, see Bond Jr and DePaulo (2006). Apart from accuracy, there was a large effect size for the AU of fear (AU 20) while differentiating lies from truth.

High-stake lies were used in some previous research. For example, Vrij and Mann (2001) used the videotaped press conferences of people who were asking for help in finding their relatives and some people were found guilty. For those materials, neither Artificial Intelligence nor a human can be sure of a veracity status or ‘ground truth’ without substantial evidence. Our database consists of high-stakes deception videos from a real game show, in which we know the veracity of the statements (there are some limits in the current game show due to the unreliable polygraph test, which can be fixed in future work using the certain ground-truth game shows such as Golden Balls, see Van den Assem et al., 2012). This kind of experimental materials has both a relatively higher ecological validity and internal validity.

Were the facial expressions in lying video clips all microexpressions (facial expressions last for from 1/25 to 1/5 of a second)? The current results of total duration showed that the average of frames of AU20 was 20.77 in truth-telling video clips and was 15.21 in lying ones, corresponding to 692ms and 507ms; the 95% confidence intervals of total duration were from 19.03 to 22.52 frames (634ms ~ 751ms) while telling truth and were from 10.32 to 20.11 frames (344ms ~ 670ms) while lying. In the current study, the mean was affected by extreme values or outliers (see Figure 1). Thus, we used the median, which could be a more appropriate statistic for the duration. The median of duration in the truth-telling video clips was 12 (400ms) and in the lying video clips was 8 (267ms). Although the duration of (partial) fear were shorter in lying video clips than in truth-telling video clips, most of the durations in lying did not fit into the limits of traditional durations of microexpressions, i.e., less than 200ms (see Shen et al. (2012). There were nearly 1/3 AU20s for which durations were less than or equal to 6 frames (200 ms) in the lying video clips, and only 1/5 of them in the truth-telling video clips were less than or equal to 6 frames. By using 500ms as the boundary between microexpressions and macroexpressions (see Matsumoto & Hwang, 2018), there were almost 2/3 of the facial expressions that could be named after microexpressions. The results suggested that the leaked emotional facial expressions in real life were much longer (the duration of apex of leaked emotional facial expressions would be less than 200ms). No matter what the duration is, or whether the facial expression is a microexpression or not, the durations of facial expressions were significantly shorter in the lying video clips than in the truth-telling video clips.

Taken together, our findings suggested that deception is detectable by using emotional facial expressions of fear in high-stake situations. Lying in the high-stake situations will leak facial expressions of fear. The durations of fear were significantly different between lying and truth-telling conditions. Besides, the facial movements will be more asymmetrical in the scenario of lying than in the scenario of telling truth.

Our findings prompted that attending to the dynamic features of AU20 (such as symmetry and duration) can improve people’s ability to differentiate liars from truth-teller. Besides, the machine learning approach may be employed to detect other real-world deceptive actions in the field of deception detection, especially those high-stake situations in which strong emotions will be generated, associated with attempts to neutral, mask, and fake such emotions (similar work is done in the project of iBorderCtrl, see Crampton, 2019).

Pupil dilation and pitch of speech are found to be significantly related to deception by some studies of meta-analysis (Bella M. DePaulo et al., 2003; Levine, 2019; Zuckerman et al., 1981). These cues are closely related to leakage too. The findings of Bradley et al. (2008) indicated that the pupil’s changes were larger when viewing emotionally arousing pictures which also were associated with increased sympathetic activity. Pitch of speech will be different between honest and deceptive interaction (Ekman et al., 1976; Zuckerman et al., 1981). Future studies should address all these leaked clues or the “hot spots” of the deception.

## 4. Materials and methods

### 4.1 The database collected by the authors

We used the video clips of the same individual who told both lies and truth in a high-stake game show. The database consisted of 32 video clips (16 persons), each individual told lies in one video clip and truth in the other.

Levine (2018) noted that cues could differ from person to person, and what spotted one liar was usually different from the signals that revealed the next liar (Levine, 2019). Meanwhile, cues may vary from sender to sender and message to message. For the same individual, however, he or she would display the almost the same pattern on different occasions. Therefore, the relatively ideal experimental materials should be composed by the same individual who tell both lies and truth to exclude the variation coming from individual differences (at least, the variation coming from the same individual should be much less than that originating from different individuals).

Considering the aforementioned variation between people and contexts, our database consists of video clips of the game show of “the moment of truth” (see https://en.wikipedia.org/wiki/The_Moment_of_Truth_(American_game_show) for details) obtained from the internet, in which the same individual tells both lies and truth. During the game show, most of the people talk emotionally because of the high-stakes situations they are in. Their emotional facial expressions are natural, rather than acting based on instructions. The ground truth was according to whether an individual was lying or not in the game show specifying by a pre-show polygraph test. Using a game show can avoid the shortcomings of real-world materials (e.g., appealing for the return of relatives) which cannot accurately be controlled over knowing the ground truth; meanwhile, the stakes in the game show can be high (the highest gain from the show can reach at 500, 000 US dollars, and cues to deception will be more pronounced than when there was no such monetary incentive, see DePaulo et al., 2003).

The video clips consist of the fragments when the individual answering the questions (from the beginning to the end of answering each question). The duration of the video clips ranges from 3 seconds to 280 seconds, with an average duration of 56.6 seconds. Because of the setting of the game show (when the individual lied the game was over), the video clips in which the individual was telling a truth were much longer than the video clips in which the individual was telling lies (105.5 s vs. 7.8 s in average, all truth-telling video fragments were merged into one video clip which duration was much longer than the lying video clip). There were 8 males and 8 females (Participants had no lies were excluded in the data set). The frame rate of all the videos was 30 f/s.

### 4.2 Using machine vision to compare the features in video clips while people lying or telling the truth

Asking people to find out the cues to deception is difficult. Furthermore, naïve human observers may not be able to perceive the subtle differences of the emotional facial expressions between telling lies and telling truth. Alternatively, machine vision may do this job well. We proposed a method aimed to use the AUs of fear to discern deceptive and honest individuals in high-stakes situations.

#### 4.2.1 Presenting the videos to a computer vision system

We used the software of OpenFace (Baltrusaitis et al., 2018) to conduct computer video analysis. The software could automatically detect the face, localize the facial landmark, output the coordination of the landmarks, and recognize the facial AUs. OpenFace can identify 18 AUs, (AU01, AU02, AU04, AU05, AU06, AU07, AU09, AU10, AU12, AU14, AU15, AU17, AU20, AU23, AU25, AU26, AU28, AU45). Furthermore, the frame-by-frame OpenFace output can give information on the intensity AUs (i.e., it can provide information on the presence and intensity of the AUs). Su and Levine (2016) showed that some AUs of emotional facial expressions can distinguish liars from truth-tellers in high-stakes situations.

According to Frank and Ekman (1997), telling a consequential lie results in emotions such as fear and guilt. Therefore, we focused on the AUs of fear, i.e., AU1, AU2, AU4, AU5, AU20, AU26.

#### 4.2.2 using MATLAB to calculate the indicators

The videos were put into OpenFace. A set of descriptors was extracted from OpenFace output frame by frame. The values of AUs of fear were generated by multiplying the output values of presence (0, 1) and the value of the intensity (from 0 to 1) for each frame; then the values of AUs of fear in each frame were aggerated and averaged (the sum of the values of AUs of fear divided by the number of frames) for further statistical analysis.

Next, we used MATLAB code to count the duration of AUs of fear (counting the number of frames when the value of the presence of corresponding AU was equal to 1). Because the frame rates of all the videos were the same, the duration of AU could be represented by the number of frames (the precise duration was obtained by dividing the total number of frames by frame rate, i.e. 30).

Beh and Goh (2019) proposed a method to detect the changes in the Euclidean distances of facial landmarks to find out microexpressions. We used the distances of ld1 and rd1, which are distances between facial landmarks at the left/right eyebrow and left/right eye (index 20/25 and index 40/43, see Figure 3), to investigate the synchronization and symmetry between left and right facial movements. The MATLAB function of Wcohenrence (wavelet coherence, the values ranged from 0 to 1) was used for this purpose, as this function returns the magnitude-squared wavelet coherence, which is a measure of the correlation between two signals (herein ld1 and rd1) in the time-frequency domain. If the left and right facial movements have perfect synchronization and symmetry, the value of wavelet coherence would be 1.

**Figure 3.**
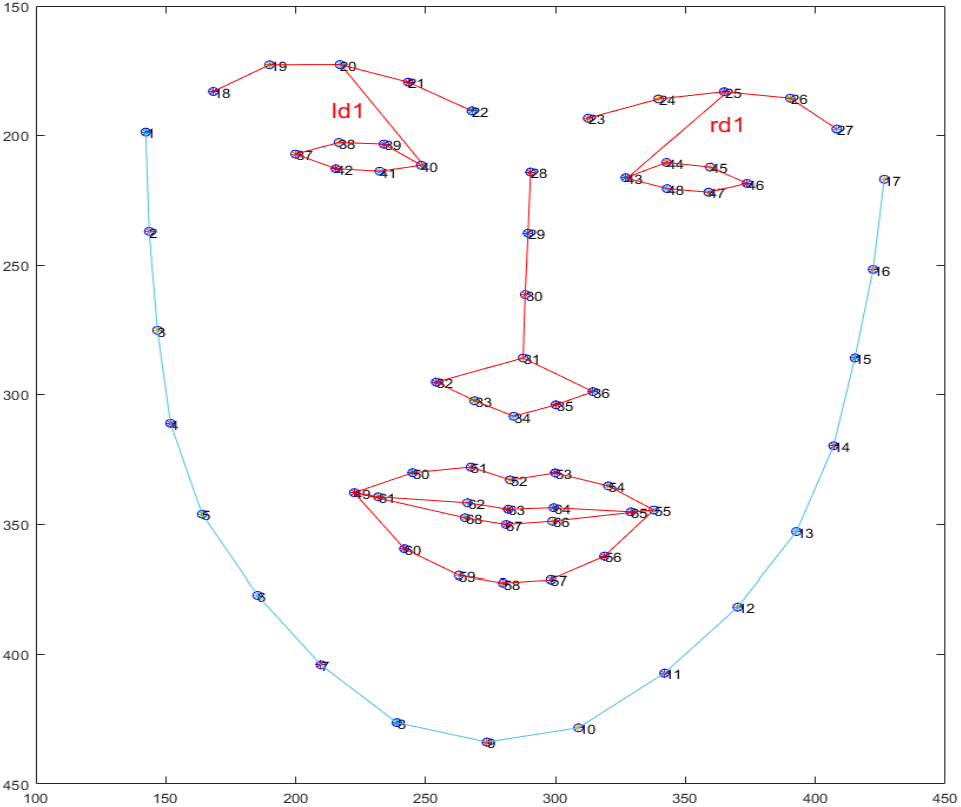
The 68 facial landmarks and the Euclidean distances of ld1 and rd1.

#### 4.2.3 using Machine Learning to classify the truth or deception

We then used WEKA(Hall et al., 2009), a Machine Learning software, to classify the videos into groups of truth and deception. Three different classifiers were trained via a 10-fold cross-validation procedure. We selected three classifiers: Random Forest, K-nearest neighbours, and Bagging. Random forest operates by constructing a multitude of decision trees which is also a better choice for data imbalance (Bruer et al., 2020). K-nearest neighbours (lazy.LBK in WEKA) achieves classification by identifying the nearest neighbours to a query example and using those neighbours to determine the class of the query (Cunningham & Delany, 2004). Bagging is a method for generating multiple versions of a predictor and using these to get an aggregated predictor (Breiman, 1996). Considering the data imbalance (the video clips of truth were much longer than the video clips of deception, 50097 frames vs. 3689 frames, which is consistent with real life that lying is not as frequent compared to truth-telling.), the data were resampled by using SMOTE (Chawla et al., 2002).

The steps of classifying the truth or deception in the video clips are demonstrated in Figure 4. First, OpenFace detected the face, localized the landmarks, output the presence and intensity of AUs. Following that, AUs of fear, as well as indicators used by Beh and Goh (2019) in each frame from both lying and truth video clips were merged into a facial movement description vector. Finally, in the classification stage, classifiers of Random Forest, K-nearest neighbours, and Bagging were trained to discriminate deception and honesty.

**Figure 4.**
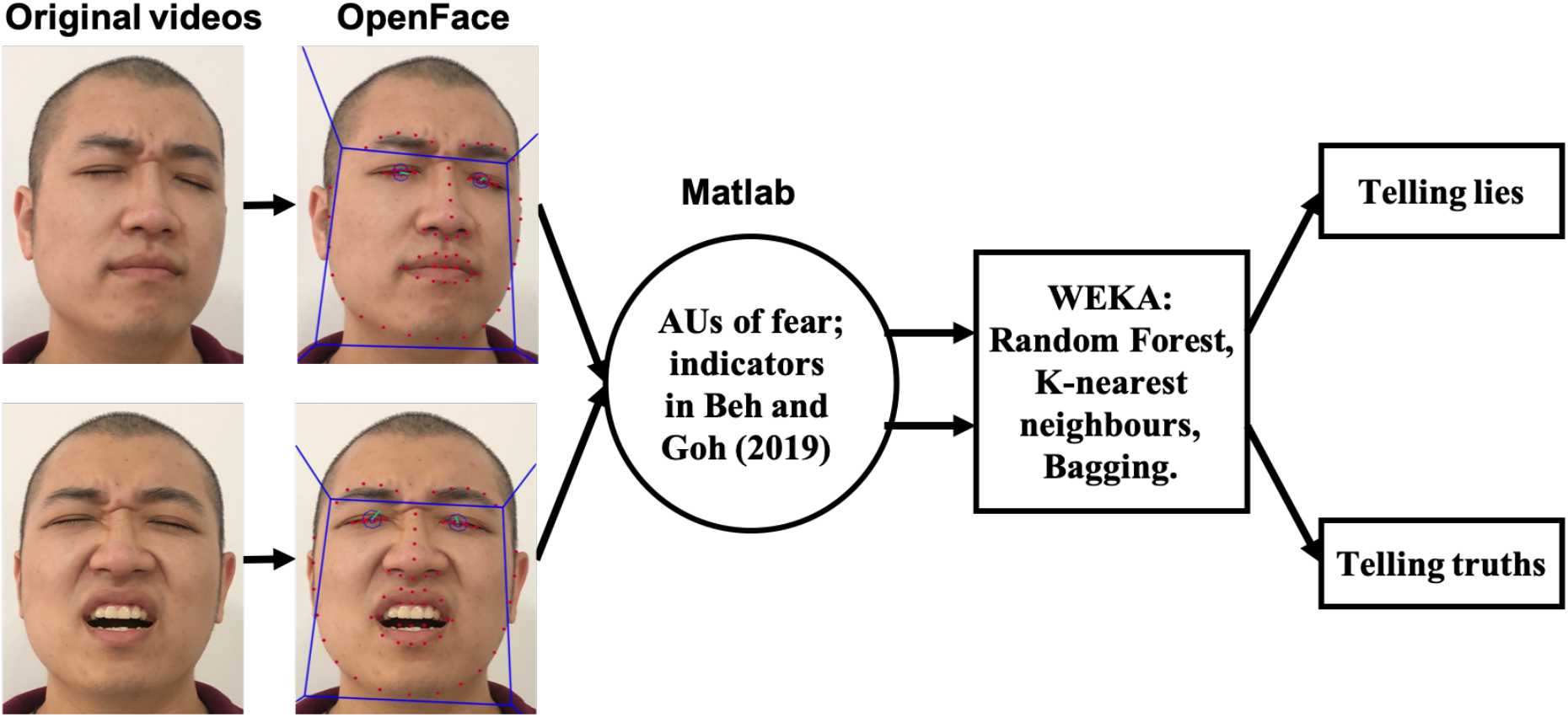
Overview of the procedure of classifying video clips. The model used here for demonstrating the processing flowchart is the third author.

## Acknowledgments

This study was partially supported by the grants from the National Natural Science Foundation of China (No. 31960180, 32000736, 31460251), the Planed Project of Social Sciences in Jiangxi Province (No. 18JY24), and the project of “1050 Young top-notch talent” of Jiangxi University of Traditional Chinese Medicine (No.5141900110, 1141900610).

